# Division of Labor: A Democratic Approach to Understanding Manual Asymmetries in Non-Human Primates

**DOI:** 10.1101/011908

**Authors:** Madhur Mangalam, Nisarg Desai, Mewa Singh

**Affiliations:** Department of Psychology, University of Georgia, Athens, United States of America; Indian Institute of Science Education and Research Pune, Pune, India; Biopsychology Laboratory, and institution of Excellence, University of Mysore, Mysore, India; Jawaharlal Nehru Center for Advanced Scientific Research, Bangalore, India

**Keywords:** capuchin monkey, *Cebus* sp, hand performance, hand preference, laterality, manual asymmetry, manual specialization

## Abstract

A consequence of the ‘gold rush’ like hunch for human-like handedness in non-human primates has been that researchers have been continually analyzing observations at the level of the population, ignoring the analysis at the level of an individual and, consequently, have potentially missed revelations on the forms and functions of manual asymmetries. Recently, consecutive studies on manual asymmetries in bonnet macaques, *Macaca radiata* [Mangalam et al., 2014a; Mangalam et al., 2014b] revealed both the functional and the adaptive significance of manual asymmetries respectively, and pointed towards the division of labor as being the general principle underlying the observed hand-usage patterns. We review the studies on manual asymmetries in capuchin monkeys, *Cebus* spp. and argue that the observed hand-usage patterns might reflect specialization of the two hands for accomplishing tasks that require different dexterity types (i.e., maneuvering in three dimensional space or physical strength). To this end, we do a step-by-step analysis of the various tasks used in the studies on manual asymmetries in capuchin monkeys, wherein we: (a) analyze the different manual tasks that have been used to study manual asymmetries in non-human primates on the basis of the attributes such as the number of hands required to solve a given task (i.e., unimanual, pseudo unimanual, or bimanual) and the spatiotemporal progression of manual actions (i.e., sequential or concurrent). (b) Determine the forms and functions of manual asymmetries that these tasks can potentially elicit within the broader scope of the behavioral repertoire of an individual, a population, or a species. (c) Qualify the scope of the inter-individual, -population, or -species comparisons. We then describe the division of labor as a general principle underlying manual asymmetries in non-human primates, and propose experimental designs that would elaborate the forms and functions of manual asymmetries in non-human primates, and the associated adaptive value.

## Introduction

Approximately 90% humans preferentially use the right hand to perform complex manual actions [Raymond and Pontier, 2004]. In order to understand the adaptive value of this population-level right-handedness, which is peculiar to humans, it is important to understand the evolutionary origin of manual asymmetries, in humans as well as in their phylogenetic relatives, the non-human primates. Manual asymmetries of some kind or the other are almost ubiquitous among the non-human primates. However, for a long time the population-level lateral bias in hand usage in non-human primates remained equivocal; considering that the exogenous factors, such as the initial position of a stimulus with respect to a subject, body posture of the subject, etc. might influence hand usage, researchers considered manual asymmetries in non-human primates to be analogous and not homologous to manual asymmetries in humans. Regardless of such an ambiguity, hand preference in non-human primates has been hypothesized to have evolved owing to functional and morphological adaptations to feeding in arboreal contexts [Bradshaw and Rogers, 1993; Papademetriou et al., 2005; Ward and Hopkins, 1993].

As opposed to the prevailing ideas on population-level right-hand preference in humans, MacNeilage et al. [1987] argued that human-like population-level lateral bias in hand usage is evident in non-human primates, and proposed the postural origins theory. According to the postural origins theory, among non-human primates initially the left hand became specialized for visually guided movements, and the right hand became specialized for postural support. Subsequently, in non-human primate species that adopted a relatively more terrestrial lifestyle, the right hand became more specialized for physical manipulation than for postural support, owing to the decreasing demands on the right hand to support vertical posture. However, the postural origins theory fails to describe why initially the left-hand (and not the right hand) became specialized for visually guided reaching, and more importantly, how a population-level right-handedness evolved during the transition from monkeys to apes to humans [McGrew and Marchant, 1997]. Overall, the postural origins theory incorporates the physical constraints on hand usage imposed by the body posture, but does not explain the variations in hand-usage patterns, corresponding to the novelty and the spatiotemporal scale of the manual actions.

In the earlier studies on manual asymmetries in non-human primates, terms such as ‘task complexity’ and ‘task demands’ were used without ever being comprehensively defined. For example, complexity of a reaching-for-food task was measured in terms of the number of steps preceding the terminal act of reaching for food, with almost no reference to the precision of movement in any of the manual actions. This made it difficult to draw any conclusions with regard to the forms and functions of manual asymmetries in non-human primates. Subsequently, based on the perspective put forward by MacNeilage et al. [1987], while simultaneously acknowledging the possibility that hand-usage patterns might vary with novelty and the spatiotemporal scale of the manual actions, as indicated by the previous studies on hand-usage patterns in non-human primates, Fagot and Vauclair [1991] put forward the task complexity theory. The task complexity theory proposes: (a) low-level tasks (i.e., tasks involving cognitively less demanding actions that are practiced frequently) elicit symmetrical hand-usage patterns at the level of the population and manual preferences at the level of an individual, not necessarily indicative of any kind of specialization. (b) High-level tasks (i.e., tasks involving cognitively more demanding manual actions that are practiced rarely) elicit asymmetrical hand-usage patterns at the level of the population, likely to be indicative of some kind of cognitive specialization. They also argued that inconsistencies in directional biases arise owing to the diversity in the tasks used to elicit manual asymmetries and the cognitive processes involved in solving them. Overall, these two types of tasks, low-level and high-level, elicit two different types of lateralization, hand preference and manual specialization.

Since the conception of the postural origins theory and the task complexity theory, there have been a plethora of studies on manual asymmetries in non-human primates with titles like “Laterality of hand functions in…,” “Hand preferences in different tasks in…,” “Consistency of hand preference across low‐level and high‐level tasks in…,” “Hand preferences in unimanual and coordinated-bimanual tasks by…,” “Posture and reaching in…,” etc. These studies generally have not independently considered the constraints consider by the task complexity theory and the postural origins theory. The task complexity theory incorporates the physical constraints imposed by tasks, whereas the postural origins theory incorporates the physical constraints imposed by body postures. These different types of physical constraints, however, may not necessarily elicit mutually consistent hand preferences. They have focused essentially on hand preference (i.e., the relative incidence of the use of either hand for responding) as the primary measure to assess manual asymmetries, with almost no reference to the forms and functions. Moreover, they have continually ignored several individual-specific traits, such as the feeding ecology and niche structure, and task-specific characteristics, such as the spatiotemporal requirements of the task, which might together influence hand-usage patterns. In such a situation, conclusions drawn from studies incorporating variable methodologies and task requirements, and not incorporating the differences between individuals, populations, or species, are likely to be misleading.

During the course our study titled “Flexibility in food extraction techniques in urban free-ranging bonnet macaques, *Macaca radiata* [Mangalam and Singh, 2013],” we observed a peculiarity in the hand-usage patterns of the study individuals. The hand used for the terminal act of reaching remained almost consistent irrespective of the number of steps involved in the food extraction process. This, rather counter-intuitive observation provoked us to carry out a systematic study on manual asymmetries in bonnet macaques. Two consecutive studies [Mangalam et al., 2014a; Mangalam et al., 2014b] revealed both the functional and the adaptive significance of manual asymmetries respectively, and pointed towards the division of labor as being the principle underlying the observed hand-usage patterns. In contrast to the conventional ideas on manual asymmetries in non‐human primates, these observations demonstrated the specialization of the two hands for tasks requiring maneuvering in three‐dimensional space or those requiring physical strength, as inferred by their consistent usage across a variety of spontaneous and experimental tasks. Also, our task apparatus revealed some peculiarities in the forms of manual asymmetries, which galvanized us to analyze the tasks used to elicit manual asymmetries in the other studies. We thus decided to summarize our analysis of these tasks and put forward our ideas on the division of labor in hand usage in the present review article.

On the basis of our studies on manual asymmetries in bonnet macaques [Mangalam et al., 2014a; Mangalam et al., 2014b], our review of studies on manual asymmetries in capuchin monkeys, *Cebus* sp., and our analysis of the various tasks used in these studies, we found that: (a) A consequence of the ‘gold rush’ like hunch for human-like handedness in non-human primates has been that researchers have been continually analyzing observations at the level of the population, ignoring the analysis at the level of an individual and, consequently, have potentially missed revelations on the forms and functions of manual asymmetries. (b) These studies lack an a priori description of a cognitively demanding and/or less-demanding manual action and the requirements of the task in terms of the form (e.g., power or precision grip; see Napier [1956]) or function (e.g., maneuvering in three-dimensional space and providing physical strength) and, therefore, remain largely contextual. (c) In multi-step tasks, even when requiring less precision, step(s) preceding the terminal act might not be a part of the behavioral repertoire of an individual, a population, or a species, in which case, inter-individual, -population, or -species comparisons of hand-usage patterns are likely to be erroneous. Thus, in the present review, we emphasize the need to explicitly study manual asymmetries in non-human primates with respect to the forms and functions, and the associated adaptive value, propose the appropriate experimental designs, and qualify the scope of inter-individual, -population, or – species comparisons.

We review the studies on manual asymmetries in capuchin monkeys, *Cebus* spp. and argue that the observed hand-usage patterns might reflect specialization of the two hands for accomplishing tasks that require different dexterity types. To this end, we do a step-by-step analysis of the various tasks used in the studies on manual asymmetries in capuchin monkeys, wherein we: (a) analyze the different manual tasks that have been used to study manual asymmetries in non-human primates on the basis of the attributes such as the number of hands required to solve a given task (i.e., unimanual, pseudo unimanual, or bimanual) and the spatiotemporal progression of manual actions (i.e., sequential or concurrent). (b) Determine the forms and functions of manual asymmetries that these tasks can potentially elicit within the broader scope of the behavioral repertoire of an individual, a population, or a species. (c) Qualify the scope of the inter-individual, -population, or -species comparisons. We then describe the division of labor as a general principle underlying manual asymmetries in non-human primates, and in order to substantiate this possibility, propose experimental designs that would elaborate the forms and functions of manual asymmetries in non-human primates, and the associated adaptive value.

## Manual Asymmetry Paradigms

Manual asymmetries did not first evolve in primates, but hemispheric specialization preceded manual symmetries instead, or in other words, evolved as a by-product of a more fundamental cerebral asymmetry affecting sensorimotor functioning [Witelson, 1988]. Accordingly, tasks that are likely to challenge the differential abilities of the two hemispheres are more likely to elicit manual asymmetries: hand preference, that is, the preferential usage of one hand to perform a unimanual task or to execute the most complex action while performing a bimanual task, or hand performance, that is, differential performance of the two hands in solving the same task [Fagot and Vauclair, 1991]. In the manual preference paradigm, repetitive presentations of a given task produce individual scores of right- and left-hand uses. These scores are then used to derive the strength and the bias of manual lateralization. The strength is obtained in several statistical ways, all of which basically calculate some index of the deviation from a random 50% hand usage regardless of the hand preferred, wherein the bias refers to the direction of manual preference (left or right). In the manual performance paradigm, on the basis of the differential reaction time or accuracy of the two hands in solving the same task individuals are classified as right- or left-handers when one hand performs better on average than the other. Studies on manual asymmetries in non-human primates make use of an array of spontaneous and experimental tasks to describe the two kinds of manual asymmetries, which we attempt analyzing below.

### (i) Quadrupedal (Pseudo) Unimanual Reaching-For-Food Tasks

Typically, quadrupedal (pseudo) unimanual reaching-for-food tasks involve reaching for food placed on the ground, on a platform, tray or in a vessel accessible directly [Fragaszy and Mitchell, 1990; Garber et al., 2008; Lilak and Phillips, 2008; Meunier and Vauclair, 2007; Parr et al., 1996; Spinozzi et al., 1998; Westergaard et al., 1997; Westergaard et al., 1998a; Westergaard and Suomi, 1993a], or through a hole [Spinozzi et al., 2004; Westergaard et al., 1998a], using one hand (here, we use the word ‘pseudo’ before unimanual because the whole process of obtaining food does involve both hands as there just cannot be any unimanual reaching-for-food task for any quadrupedal individual).

An appropriate assessment of hand preference with regard to unimanual reaching-for-food tasks has several underlying assumptions: (a) a subject is equally likely to use any of its two hands, which is practically possible only when the subject is acquiring either sitting or bipedal posture such that there are no ergonomic constraints on the usage of any of the two hands. (b) Food is located exactly on the sagittal plane of the body of the subject so that its spatial arrangement does not influence hand preference (though this assumption is almost always met as there is an equal probability of food being located towards the right and left of the sagittal plane).

Whereas quadrupedal (pseudo) unimanual reaching-for-food tasks are assumed to involve only one hand, they implicitly involve the other hand which is required to passively maintain tripedal posture. This hand faces an increase in physical load when the other hand is set free for prehension. Thus, one hand is used to maintain tripedal posture and the other hand is used to maneuver in three-dimensional space or to make precision grips, following the principle of division of labor. Also, under experimental conditions, ergonomic constraints imposed by the possible asymmetries in the body posture of an individual, together with or independent of the preferential use of one hand for maintaining tripedal posture, is likely to influence hand preference in quadrupedal (pseudo) unimanual reaching-for-food tasks. However, studies on hand preference in capuchins have drawn conclusions with regard to the effect of the complexity of the tasks on hand preference without ever deploying a purely unimanual task independent of these influences.

### (ii) Bipedal (Pseudo) Unimanual Reaching-For-Food Task

Typically, bipedal (pseudo) unimanual reaching-for-food tasks involve obtaining a single piece of food placed on a high-rise platform, tray or in a vessel accessible directly [Spinozzi et al., 1998; Westergaard et al., 1997; Westergaard et al., 1998a] or through a hole [Parr et al., 1996; Westergaard et al., 1998a], using one hand (as in the case of the quadrupedal (pseudo) unimanual reaching-for-food tasks, we use the word ‘pseudo’ before unimanual).

Bipedal (pseudo) unimanual reaching-for-food tasks can only be solved using both hands and in no less than two or three steps: (P1) two-step process: step 1: setting one hand, hand-1 (i.e., either left or right hand), free from maintaining quadrupedal posture and using it to hold a high-rise structure (this action is physically demanding as the body is liXfted/pulled upwards) while maintaining tripedal posture using the other hand, hand-2; step 2: setting the other hand, hand-2, free from tripedal posture and using it to reach for food while maintaining bipedal posture using the other hand, hand-1. (P2) Three-step process: step 1: setting one hand, hand-1, free from maintaining quadrupedal posture and using it to hold a high-rise structure (as mentioned above, this action is physically demanding as the body is lifted/pulled upwards) while maintaining tripedal posture using the other hand, hand-2; step 2: setting the other hand, hand-1, free from tripedal posture and using it to hold the high-rise structure; step 3: using one hand, (P1a) hand-1 (in which case the sequence is functionally similar to the previous one) or (P2b) hand-2, to reach for food.

These sequences of manual actions involve both hands, following the principle of division of labor, that is, one hand is used to perform the actions demanding relatively more physical strength (e.g., lifting/pulling the body) and the other hand is used to perform the actions demanding more sophistication (e.g., making precision grips or maneuvering in three-dimensional space). However, studies on hand preference in capuchins have almost never reported the stepwise usage of the two hands for solving bipedal (pseudo) unimanual reaching-for-for-food tasks as described above, restricting their data collection and analysis only to manual actions that are directly associated with prehension. Comparative assessment of hand preference in the quadrupedal and bipedal (pseudo) unimanual reaching-for-food tasks, as reported by Spinozzi et al. [1998] and Westergaard et al. [1997, 1998], demonstrates that capuchins consistently use one hand for prehension in both types of tasks, which is possible only while following either the two-step process (i.e., P1) or the second of the three-step processes (i.e., P2b) for solving bipedal (pseudo) unimanual reaching-for-food tasks.

### (iii) Quadrupedal/Bipedal Coordinated Bimanual Task

Typically, solving a coordinated bimanual task involves obtaining food from ~ 10 to 15 cm long and ~ 3 to 5 cm wide transparent/opaque tube [Lilak and Phillips, 2008; Meunier and Vauclair, 2007; Spinozzi et al., 1998; Spinozzi et al., 2007; Westergaard and Suomi, 1998]. An individual that is assuming a quadrupedal position can solve the task in two or three steps: (P1) step 1: picking up the tube with one hand, hand-1, while maintaining tripedal posture with the other hand, hand-2; step 2: attaining bipedal posture by freeing hand-2 and extracting the food from the tube with the same hand. (P2) step 1: picking up the tube with one hand, hand-1, while maintaining tripedal posture with the other hand, hand-2; step 2: attaining bipedal posture by freeing hand-2, and shifting the tube from hand-1 to hand-2; step 3: extracting the food with hand-1. Thus, it needs to be determined whether an individual continued holding the tube with the same hand or shifted it to the other hand. In case of the shift the observed hand-usage pattern can be explained using the principle of the division of labor (as described by Mangalam et al. [2014a]); and in the other case as well as when an individual is assuming a bipedal posture while picking up the tube, sequential planning of motor actions. However, studies do not analyze manual asymmetries in solving coordinated bimanual tube task from this perspective and, therefore, present only a partial picture.

### (iv) Sequential Unimanual/Bimanual versus Concurrent Bimanual Tasks

Typically, solving a box task involves obtaining a single piece of food placed on a tray inside a clear plexiglass box. The box can be opened by lifting its lid that is hinged to one of its walls. There are two different versions of the box task. In one version, the lid may remain open once it is lifted beyond a point [Lilak and Phillips, 2008; Spinozzi and Truppa, 2002], in which case the task can be solved in either 2 steps: lifting the lid and reaching for food, in a sequential unimanual/bimanual manner (L-L/R-R, L-R/R-L, B-L/B-R); or 3 steps: lifting the lid, holding the lid up, and reaching for food, in a concurrent bimanual manner (L-RL/R-LR, L-LR/R-RL, B-LR/B-RL). In another version, the box includes a stop screw on the back of the lid which causes the lid to fall shut if it is not held open [Lilak and Phillips, 2008; Spinozzi and Truppa, 2002], in which case the task can be solved only in 3 steps: lifting the lid, holding the lid up, and reaching for food, in a concurrent bimanual manner (L-RL/R-LR, L-LR/R-RL, B-LR/B-RL; in the latter two cases, the sequence is functionally similar to the previous one).

Spinozzi and Truppa [2002] did an assessment of hand preference in 23 tufted capuchins using the box tasks. While solving the sequential unimanual/bimanual box task, the capuchins indiscriminately (in 48.8% and 36.9% trials) used the strategies involving no differentiation (L-L/R-R, i.e., lifting the lid and reaching for food with the same hand), and differentiation of roles for the two hands (L-R/R-L, i.e., lifting the lid with one hand and reaching for food with the other hand); and while solving the concurrent bimanual version of the task, the capuchins predominantly (in 73.4% trials) used the strategy involving complete differentiation of roles for the two hands (L-LR/R-RL, i.e., lifting the lid and holding it up with the same hand, while simultaneously reaching for food with the other hand) more often than the other two possible strategies (L-RL/R-LR and B-LR/B-RL). In a nutshell, the capuchins did not show any difference in the direction and strength of hand preference for prehension between the sequential unimanual/bimanual and concurrent bimanual versions of the box task, demonstrating the similarity between them.

This example demonstrates that sequential unimanual/bimanual and concurrent bimanual box tasks elicit similar direction and strength of hand preference. This also holds true for several other tasks as described above. In fact, a general principle involving partial/complete differentiation of roles for the two hands is likely to underlie manual asymmetries and, therefore, sequential unimanual/bimanual and concurrent bimanual tasks should not be treated differently.

### (v) Haptic Search Tasks

Typically, solving a haptic search task involves obtaining food mixed with some non-edible material [Parr et al., 1996; Spinozzi and Cacchiarelli, 2000] or placed in the crevices on the surface of variably shaped objects [Lacreuse, 1999; Lacreuse and Fragaszy, 1996; Lacreuse and Fragaszy, 1997] from the inside of an opaque box (~ 15 to 30 cm × 15 to 30 cm × 15 to 30 cm) through a small opening (diameter < 5 cm; these dimensions allow inserting only one hand at a time). Haptic discrimination has been found to be more difficult that visual discrimination in non-human primates (see, for example, Wilson [1965] in rhesus macaques), perhaps because haptic perception without visual guidance is uncommon in natural settings. Thus, haptic judgments are likely to be novel and consequently, cognitively more demanding as compared to visually guided judgments. Studies on manual asymmetries therefore make use of haptic search tasks to differentially challenge the perceptual motor abilities of the hands, which are likely to be affected by functional differences between the left and right hemispheres. However, studies do not compare hand-usage patterns between haptic and visually guided reaching (though Spinozzi and Cacchiarelli [2000] and Lacreuse [1999] stand out as an exception), rather just describe manual asymmetries in haptic search tasks; this hardy reveals something substantial as studying haptic judgments in isolation from visually guided judgments, fail to resolve manual asymmetries stemming from the absence of the visual cues alone.

### (vi) Probing/Tool-Using Tasks

Typically, solving a (pseudo) unimanual probing task involves manipulating a wooden dowel inserted into a small hole in a clear Plexiglas box in order to displace a food reward off a shelf where it could be retrieved manually [Garber et al., 2008], using a stick to obtain food material present inside a vessel with a narrow opening while maintaining a tripedal posture [Anderson et al., 1996; Westergaard et al., 1998a; Westergaard et al., 1998b; Westergaard and Suomi, 1994a; Westergaard and Suomi, 1994b] (another version may involve using a sponge [Westergaard and Suomi, 1993a]) or a bipedal posture [Lilak and Phillips, 2008; Westergaard, 1991; Westergaard et al., 1998a]; another tool-using task is nut-cracking that involves coordinated bimanual handling of stones to crack nuts [Westergaard and Suomi, 1993b; Westergaard and Suomi, 1996]. It is important to note here that the above probing/tool-using tasks are similar in terms of the number of hands required to solve the task (i.e., unimanual, pseudo unimanual, or bimanual) and the spatiotemporal progression of manual actions (i.e., sequential or concurrent) except for the fact that they involve an extension of the body, controlling which requires finer finger adjustments through response-produced feedback. Thus, functionally similar to simple reaching-for-food tasks, probing/tool-using tasks are likely to prove helpful only if the form of manual asymmetries (i.e., with respect to grip type) is considered.

### (vii) Spontaneous Tasks

Hand-usage patterns in tasks such as grooming [Fragaszy and Mitchell, 1990], maternal cradling and infant positioning [Hopkins, 2004; Panger and Wolfe, 2000; Westergaard et al., 1999]are more likely to be influenced by the specialization of the two hands for more common activities such as feeding than these tasks themselves. For example, a female capuchin which has its left hand specialized for fine finer adjustments or maneuvering in three dimensional space and its right hand specialized for physical support is more likely to use its right hand for maternal cradling and infant positioning just to keep its left hand free for the usual feeding activities (as they require more sophisticated manual actions). However, studies merely describe the hand used for these activities without considering the forms and functions of the associated manual asymmetries.

## Forms and Functions of Manual Asymmetries

The corticomotoneuronal connections innervating the hands regulate the timing and precision of the muscular forces required for fine finger adjustments through response-produced feedback (see, for example, Porter [1985]). It follows from this fact that actions with finer sequential finger movements are more likely to elicit manual asymmetries than simpler actions, as Elliott and Chua [1996] proposed in humans (also see Healey et al. [1986], Steenhuis [1996], and Steenhuis and Bryden [1989]). There exists a possibility that lateral asymmetry in the number of corticomotoneuronal connections innervating the hands govern the forms and functions of manual asymmetries: the hand with lesser corticomotoneuronal connections is specialized for manual operations that primarily involve physical strength or those that require power grips, and the hand with greater corticomotoneuronal connections is specialized for manual actions that involve maneuvering in three-dimensional space or those that require precision grips (see Mangalam et al. [2014b]). A step-by-step analysis of any of the above tasks reveals sequential or concurrent fundamental manual actions. These fundamental manual actions can then be classified in terms of the form into either the power or precision grip, or in terms of the function into either ‘maneuvering in three dimensional space or providing physical strength.

## Inter-Individual, -Population, or -Species Comparisons

Some intermediate step(s) involved in solving a multi-step task might not be a part of the behavioral repertoire of an individual, a population, or a species. Consequently, the perceived complexity of a task might vary across individuals, populations, or species, making inter-individual, -population, or -species comparisons of hand preferences across complex tasks erroneous. Diversity in factors causing spatiotemporal inter-individual, -population, or -species variations in manual actions may also influence hand-usage patterns at multiple levels of organization. For example, Sfar et al. [2014] did a comparative assessment of hand preference in red howlers, *Alouatta seniculus* and yellow-breasted capuchins, *Sapajus xanthosternos*: the red howlers, which habitually use the mouth to obtain food, selectively took part in the reaching-for-food tasks and also exhibited stronger hand preferences than the yellow-breasted capuchins in the tasks that were relatively simple to solve. However, differences in the strength of hand preference diminished with the increasing complexity of the reaching-for-food tasks, that is, the relatively more complex tasks were perceived as equally complex by both the red howlers and the yellow-breasted capuchins. Both these observations demonstrate that the feeding ecology and niche structure influence hand-usage patterns, bringing about the differences in hand preference out of the contingent nature of the complexity of a task. Thus, manual asymmetries in non-human primates should be investigated not just in isolation, but within the broader scope of the behavioral repertoire of an individual, a population, or a species.

## Division of Labor as a General Principle

Our experience with studies on hand-usage patterns in bonnet macaques [Mangalam et al., 2014a; Mangalam et al., 2014b; Sfar et al., 2014], our review of studies on hand-usage patterns in capuchins, and our analysis of various tasks used in these studies, collectively suggest that ‘division of labor’ is a general principle underlying manual asymmetries in non-human primates. In order to substantiate this possibility, we propose that:

### (i) Division of Labor in Hand Usage Is Likely to Be Prominently Visible in Transitions between Tasks with Variable Requirements

Individuals may have to make transitions between tasks with variable requirements and depending on these, vary hand usage. Suppose, for example, an individual that preferentially uses the left hand to make power grips and the right hand to make precision grips is solving a reaching-for-food task that involves obtaining food items from a portable container (e.g., a water bottle); the individual holds the container in the left hand and retrieves the food items with the right hand. A conspecific then approaches this focal individual and so it moves with the bottle to some other location, say to a nearby high-rise platform, or to a distant branch. There can be two ways an individual can do that: (a) by holding the bottle in the left hand and climbing with the right hand or (b) by shifting the bottle to the right hand, setting the left hand free, and climbing with the left hand. If one hand is specialized for manual operations that require power grips and the other hand is specialized for manual operations that require precision grips, or alternatively for maneuvering in three-dimensional space and providing physical strength, the second way seems more plausible (see Mangalam et al. [2014b] for another such example). So, if the transition involves tasks with variable requirements, division of labor becomes evident. We propose an experimental design to observe the division of labor in hand usage based on task demands. One should examine hand preference across situations synonymous to that in the above example. Stringent changes in hand-usage patterns while shifting contexts would demonstrate division of labor in hand usage.

### (ii) Division of Labor in Hand Usage Is Likely to Be Visible and Understood in Tasks with Differential Requirements

Napier [1956] described prehensile functions of the human hand, such as grasping and gripping: an object can be grasped/gripped by either holding it in a clamp formed by partly flexed fingers and palm, while applying a counter pressure by the thumb lying more or less in plane of the palm–the ‘power’ grip, or pinching it between the flexor aspects of the fingers and the opposing thumb–the ‘precision’ grip. Performing certain manual operations primarily requires power and precision plays a secondary role, whereas performing certain other manual operations primarily requires precision and power plays a secondary role. And this task-specific requirement of power and precision grip is likely to influence hand-usage patterns in a given manual operation. In New World monkey species, the typical hinge-shaped joint of the thumb at the base of the palm allows abduction/adduction and flexion/extension movements, but not rotational movement, the key factor in opposability [Napier and Napier, 1967]. For a long time it was thus held, that no New World monkey species could grasp objects with precision [Bishop, 1964; Napier, 1993; Napier and Napier, 1967]. However, comparative behavioral studies demonstrated that capuchins stand out from other platyrrhine species because of their (a) high degree of manual dexterity [Fragaszy, 1986; Lacreuse and Fragaszy, 1996; Panger, 1988], (b) frequent use of precision grips that mainly involve lateral aspects of digits for picking up small objects [Christel and Fragaszy, 2000; Costello and Fragaszy, 1988; Spinozzi et al., 2004], and (c) capacity to perform relatively independent movements of the digits [Christel and Fragaszy, 2000; Costello and Fragaszy, 1988].

Anatomical and physiological features of the neural substrate that control manual actions might explain the high manual dexterity in capuchins. Capuchins can act out highly fractionated movements of the fingers/digits owing to the large number and extension of the corticomotoneuronal connections that innervate the hand [Kuypers, 1981; Lemon, 1993; Muir and Lemon, 1983; Shinoda et al., 1981], as observed in humans and chimpanzees [Bortoff and Strick, 1993]. Moreover, studies reported that the individuals that preferentially used the right hand to reach for food in a concurrent bimanual tube task, exhibited a greater leftward bias of the anterior cerebellum [Phillips and Hopkins, 2007], and had a shallower central sulcus [Phillips and Sherwood, 2005] as well as a smaller overall corpus callosum in the contralateral hemisphere [Phillips et al., 2007], compared to those that preferentially used the left hand or did not show hand preference; although there was no difference in the size of the left-frontal petalia between the two [Phillips and Sherwood, 2007].

A few studies investigated manual asymmetries with respect to the control and movement of the fingers/digits in capuchins. Christel and Fragaszy [2000] reported that the individuals did not exhibit considerable patterns in hand preference or hand performance with respect to the power or precision grips used to grasp currants and grapes lying on a tray. Spinozzi et al. [2004] reported that the individuals preferentially used one hand to grasp a food item fixed on a tray, and did not show any difference in performance with respect to the power or precision grips, but extracted the food faster with the preferred hand than the non-preferred hand with respect to the precision grips (and not with respect to the power grips). Spinozzi et al. [2007] reported that the individuals preferentially used one hand to retrieve a raisin from a transparent hollow tube fixed horizontally to the upper end of a vertical metal bar, and extracted the food faster with the preferred hand than the other hand. Whereas these findings indicate that precise control/movement of the fingers/digits are more likely to elicit manual asymmetries than the imprecise ones, there are problems with the experimental setups.

If, suppose, one hand is specialized for manual operations that primarily involve physical strength and, therefore, require power grips, and the other hand is specialized for those that involve maneuvering in three-dimensional space and, therefore, require precision grips, a manual operation that primarily requires either one or the other of the two forms and functions of the hand is likely to influence hand-usage patterns with respect to a particular type of grip as well as grip-formation patterns with respect to a particular hand. The three studies–Christel and Fragaszy [2000], Spinozzi et al. [2004], and Spinozzi et al. [2007]–employ reaching-for-food tasks that primarily involve maneuvering in three-dimensional space and, therefore, require precision grip. This is likely to be the reason why Christel and Fragaszy [2000] did not find manual asymmetries with respect to the types of grips and Spinozzi et al. [2004] did not find a difference in performance between the two hands with respect to the power grips, presenting a distorted and partial picture of manual asymmetries.

We propose an experimental design to unambiguously determining the forms and functions of manual asymmetries in non-human primates. One should examine hand preference in a concurrent, bimanual reaching-for-food task. In one scenario, the manual operations should require a power grip followed by a precision grip; in another scenario, the manual operations should require a precision grip followed by a power grip. Contrasting hand-usage patterns in these two scenarios would indicate that the individuals preferentially used the two hands depending on the requirements of the tasks, that is, one hand to perform the manual operations involving maneuvering in three-dimensional space and the other hand to perform those involving physical strength. One should then examine hand performance with regard to the requirements of the tasks in a concurrent, bimanual hand-performance-differentiation task. In one scenario, this task should ergonomically force the usage of either the left or the right hand to perform a manual operation requiring either a power grip or a precision grip; in another scenario, this task should ergonomically force the usage of either the left or the right hand to perform a manual operation requiring a precision grip and the other hand to perform the one requiring a power grip. A more effective and/or efficient power grip in one scenario and a precision grip in the other scenario would indicate that the individuals used the two hands depending on the specializations, that is, difference in the manual dexterity of the two hands.

### (iii) Division of Labor in Hand Usage Is Likely to Improve Hand Performance in Terms of the Efficiency of the Power and Precision Grips

Manual asymmetries might have ecological disadvantages as they can potentially make an individual vulnerable to attack/defend appropriately only when the prey/predator is present on a particular side. Also, as the stimuli are randomly located with respect to the sagittal plane of an individual, i.e., towards left or towards right, it might make it difficult to solve a particular task. However, manual asymmetries are likely to help increasing manual specialization, the benefits of which surpass the associated ecological disadvantages (reviewed by Vallortigara and Rogers [2005]). Trehub [1983] drew a distinction between mere hand preference and manual specialization by exemplifying human infants who exhibit manual specialization and not hand preference (this idea was carried forward by Fagot and Vauclair [1991] in non-human primates). According to Trehub [1983], hand preference refers to the consistent usage of one hand to solve familiar, relatively simple, and highly practiced tasks, and may not be necessarily accompanied by an improvement in hand performance; whereas manual specialization refers to the consistent usage of one hand to solve novel, relatively complex, and not-practiced tasks that require peculiar action patterns, and is necessarily accompanied by an improvement in hand performance. Trehub [1983] also described that individuals generally exhibit manual specialization only in the context of tasks that involve cognitively demanding manual actions (see, for example, Mangalam et al. [2014b] that showed manual specialization in bonnet macaques in tasks requiring peculiar action patterns viz., in terms of tasks that require either higher maneuvering dexterity or higher physical strength). Thus, there exists a marked difference between hand preference and manual specialization in terms of the resulting difference in performance of the two hands, evidently visible while considering the forms and functions of manual asymmetries, as described in the previous section.

Only one study examined the relationship between strength of hand preference and the corresponding hand performance in capuchins. Fragaszy and Mitchell [1990] reported that the individuals exhibited a weak, but statistically non-significant, positive relationship between strength of hand preference and the corresponding hand performance in the (pseudo) unimanual and bimanual versions of the box task. However, Fragaszy and Mitchell [1990] acknowledged that the strength of hand preference could have affected the timing of the hand movements, thereby affecting the relationship between strength of hand preference and the corresponding hand performance. A similar study in another non-human primate species–the bonnet macaque, Mangalam et al. [2014a], reported a negative relationship between (a) hand performance of the preferred hand and the difference in hand performance between the two hands in a hand-performance-differentiation task, and (b) difference in hand performance between the two hands and the difference in the strength of hand preference in another (pseudo) unimanual and bimanual versions of the box task in bonnet macaques. These findings indicate that a greater strength of hand preference is associated with a higher difference in the performance of the two hands. However, research lacks sufficient evidence supporting the hypothesis that hand preference, or better yet, division of labor in hand usage improves hand performance in terms of the time and/or energy required to perform a given task.

We propose an experimental design to determine the adaptive value of hand preference. One should examine hand preference in a (pseudo) unimanual reaching-for-food task (wherein, the manual operation should require either a power grip or a precision grip) and a concurrent, bimanual reaching-for-food task (wherein, the manual operations should require a power grip with one hand followed by a precision grip with the other hand, or a precision grip with one hand followed by a power grip with the other hand). One should then examine hand performance in a hand-performance-differentiation task that should ergonomically force the usage of either the left or the right hand to perform a manual operation requiring either a power grip or a precision grip, thus allowing to measure hand performance independent of ceiling effects as this task is unlikely to elicit, or better yet, prime any motor actions associated with the opposite hand). A positive relationship between (a) hand performance of the hand with higher performance in the hand-performance-differentiation task and normalized difference in hand performance for the two hands, and (b) difference in hand performance for the two hands in the hand-performance-differentiation task and difference in strength of hand preference in the (pseudo) unimanual and bimanual reaching-for-food tasks, with respect to the power grips, the precision grips, or both, would indicate that the division of labor in hand usage improves hand performance.

## Conclusions

Studies have investigated the evolutionary origin of hand-preference in non-human primates. A careful analysis points towards the division of labor as being a general principle underlying manual asymmetries. This principle is based on the difference in the intrinsic requirements of the tasks, which can be broadly divided into maneuvering in three-dimensional space and providing physical support, acquiring power and precision grips respectively. Our review of studies on hand-usage patterns in non-human primates reveals conceptual and logistic problems with the spontaneous and experimental tasks used to determine hand-usage patterns; moreover, methodology differs and confounding variables are often not appropriately addressed. We suggest that studies on manual asymmetries in non-human primates should design experiments that do not undermine this possibility. As far as the adaptive value of manual asymmetries are concerned, we suggest that, to obtain more unambiguous answers, studies should be conducted with experimental designs that allow comparing hand-usage patterns across species that vary in their phylogenetic relatedness and/or ecology, over a range of spontaneous activities and experimental tasks. It might be useful to study manual preferences not just in isolation, but within the broader scope of the behavioral repertoire of the species. Also, it might be advantageous to study the ontogeny of manual preferences. Studies of these kinds may help to understand the forms and functions of manual asymmetries, and the potential selection pressures under which manual asymmetries are likely to appear and evolve.

